# “Cell-in-Water” measurements using Optical Photothermal Infrared spectroscopy

**DOI:** 10.64898/2025.12.28.696748

**Authors:** Wiktoria Korona, Anna Maria Nowakowska, Anna Pieczara, Barbara Orzechowska, Miriam Unger, Malgorzata Baranska

## Abstract

Conventional Fourier transform infrared absorption spectroscopy (FT-IR) allows for measurements of spatial resolution comparable to a whole cell size, which are performed on dried cells to avoid water interference. However, such conditions can disrupt the native morphology and chemical composition of cells, whilst not providing for crucial sub-cellular biochemical detail. Pump-probe Optical Photothermal Infrared (O-PTIR) Microscopy offers high-resolution measurements of cells, but the challenge of measuring cells without disturbing their morphology remains. To overcome this, we developed a protocol enabling O-PTIR measurements of cells in aqueous environments using a silicone-sealed microchamber, which allows imaging of live cells in 2 µl of buffer for up to 4 hours. Analysis of O-PTIR data obtained from hydrated and dehydrated cells confirms that hydrated conditions better preserve subcellular organization, as evidenced by secondary structure changes of proteins. We also propose a step-by-step algorithm that can improve image quality through ratiometric analysis, histogram assessment, and the application of RGB overlays. This approach allows for visualization of multiple subcellular components, such as the nucleus or lipid droplets. Trough the combination of “sandwich” sample configuration and O-PTIR, we offer a promising strategy to advance O-PTIR measurements of cells in water, which have long been considered a major challenge. A direct comparison of O-PTIR with Raman microscopy highlights the complementarity of these two techniques and similar spatial resolution.

## 1. Introduction

Non-invasive cell imaging techniques are used to study and monitor cellular dysfunctions associated with many lifestyle diseases^1^. Such studies can be performed at the molecular level and can be a part of metabolomics, i.e. comprehensive analysis of endogenous small molecules associated with specific metabolic pathways^2^. A key challenge is to capture the complexity of biochemical processes in their native state. Typically, metabolomics protocols are based on mass spectrometry and require the isolation of metabolites, which is carried out at multiple steps^3^, so it affects the integrity and composition of the sample.

Spectroscopic techniques such as Raman spectroscopy (RS), Fourier-transform infrared spectroscopy (FT-IR), and optical photothermal infrared (O-PTIR) microscopy can be used to study cells in a nondestructive way. Cell imaging provides detailed insights into cellular structures, dynamics, and biochemical composition^5–8^. To some extent, vibrational spectroscopic techniques were used for monitoring of cellular processes and quantitative analysis, making them highly valuable in biomedical research^9,10^. ***Table 1*** presents a comparison of RS, FT-IR, and O-PTIR spectroscopy in the context of their applications for cellular imaging. They are complementary, providing information on the complex biochemical composition of cells.

**Table 1.** Comparison of properties of RS, FT-IR, and O-PTIR in cell imaging^14,15^.

| feature | RS | FT-IR | O-PTIR |
| --- | --- | --- | --- |
| speed | slow | fast | fast |
| spatial resolution | high ( ~300 nm) | low (~1–10 μm) | high (~300 nm) |
| signal generation | polarizability | dipole moment | thermal expansion |
| water | not a problem | an interferent | much less of an artefact |
| live cell imaging | possible | very challenging | challenging |
| SNR | weak | relatively high | high |

Pump-probe O-PTIR imaging offers up to a 40-fold improvement in spatial resolution compared to conventional FT-IR, allowing for subcellular chemical analysis^11^. Furthermore, time-resolved studies demonstrated that cytoplasm, nucleus, and lipid droplets (LDs) exhibit distinguishable properties enabling the differentiation of subtle biomolecular signals from the dominant water background, typically observed in IR spectroscopy^12,13^. In addition, O-PTIR enables single-channel imaging, in which contrast is generated by selectively probing a chosen vibrational mode. This targeted approach allows specific chemical functionalities to be highlighted, thereby simplifying spectral interpretation and enhancing sensitivity to selected biomolecular features. This feature offers clear practical advantages for high-throughput analyses of predefined specific molecular markers and for experiments employing spectroscopic labels.

RS can be used to analyze both dried and hydrated samples under physiological conditions, with optimal performance in aqueous environments. However, IR-based techniques are gaining more and more interest in studying biological systems due to larger cross sections for absorption than noncoherent Raman scattering, which enables the detection of biomolecules even at low concentrations^16,17^. So far, most FT-IR and O-PTIR applications have included analysis of the chemical composition of dried samples due to the considerable influence of signals from water, which, to some extent, affected biologically relevant information (especially detected in the amide I region of the IR spectrum). The O-PTIR technique requires the usage of the pump-probe strategy with visible light to detect IR absorption by tracking thermal expansion and refractive index changes in the sample. This approach not only breaks the diffraction limit of conventional IR spectroscopy (limited to a few µm) but also significantly reduces water interference^16,17^. In O-PTIR technique, the IR absorption is measured indirectly through changes in the sample’s density and refractive index caused by local heating. Due to the high heat capacity of water, the increase in water temperature during O-PTIR measurement is minimal, which translates into reduced water background in O-PTIR images in comparison to the FT-IR ones. This advantage makes O-PTIR an up-and-coming technique for studying biological systems, including living cells in aqueous environments. In addition, various measurement configurations of the detection method are being developed to further reduce the water background, such as lock-in free photothermal dynamic imaging^13,16^, boxar detection^18^, as well as data analysis protocols^19^.

Recent studies have demonstrated that O-PTIR can be successfully applied for the spectral characterization of cells in the buffer. However, choosing an effective sample preparation method and measurement configuration has proven challenging, but is possible. In 2016, Cheng *et al*. pioneered in live cell O-PTIR imaging using a home-built system and designed CaF_2_ Petri dishes^20^. They achieved submicrometer-resolution in the O-PTIR imaging of lipid droplets in living prostate cancer cells or cellular drug uptake^20^. In 2021 Gardner *et al*. compared O-PTIR and Raman spectra of live and fixed cancer cells collected from the same location, providing a comprehensive characterization of the molecular structure of cells. For live cell measurements they used a “sandwich” configuration built of CaF_2_ windows (with a thickness of 500 µm) and a peroxidase−antiperoxidase pen, which was used to form a thin hydrophobic film on the edge of the window to seal the constructed chamber hermetically^21^. Fujita *et al*. performed O-PTIR imaging of trifluoromethoxy carbonyl cyanide phenylhydrazone (FCCP) in living HeLa cells using a home-built photothermal microscopy system^22^ and a “sandwich” configuration composed of two CaF_2_ cover slips (with a thickness of 170 μm) with the use of a silicone spacer (with a thickness of 100 μm). In this configuration, a signal-to-noise ratio of O-PTIR images of the nitrile group proved to be several tens of times better than for stimulated Raman imaging. A simpler configuration with two CaF_2_ windows (with a thickness of 200 µm) without the spacer was successfully used by Krafft *et al*. in 2023 to measure various cancer cells in an aqueous environment^23^. Ideguchi *et al*. demonstrated a high-temporal resolution O-PTIR imaging of live cells using a homemade wide-field quantitative phase microscope^24^. They also used a “sandwich” configuration with CaF_2_ windows of the thickness of 500 µm and showed tracking H_2_O/D_2_O exchange in live cells through aquaporins.

Studies of cell metabolism under physiological conditions have focused primarily on lipids, e.g. *de novo* lipogenesis in adipocytes using isotopically labeled glucose (^13^C)^17^, effect of oleic acid uptake of *de novo* synthesized lipids storage in LDs in hepatocytes and adipocytes^25^, or neutral lipids in LDs of living human cancer cells traced in real time using ^2^H-labeled palmitic acid^26^. A widely adopted approach in O-PTIR-based live-cell studies involved cell labeling, which enhances molecular specificity and enables dynamic tracking of metabolic processes with minimal perturbation. Cheng *et al*. developed various vibrational probes to monitor transformations in metabolic pathways of living cells, including enzymatic activity^27^ and carbohydrate trafficking^19^, even in real time. He *et al*. introduced nitrile-tagged enzyme activity labels, known as nitrile chameleons, which exhibit peak position shifts upon conversion from substrate to product. This strategy enables direct monitoring of enzymatic reactions and allows qualitative imaging of enzyme activity distribution at a spatial resolution of 300 nm^27^. These cutting-edge applications highlight the expanding capabilities of the O-PTIR technique in live-cell analysis.

In comparison to the previous studies, this work aims to develop a novel analytical approach, including sample preparation, measurement conditions, and data analysis, that enables the measurement and analysis of cells under physiological conditions using the O-PTIR technique. First, the optimal method for sample preparation was developed, and then, the collected data, primarily the single-channel O-PTIR images, were analyzed. Despite previously reported solutions, O-PTIR microscopy of cells in water has remained constrained by both instrumental and analytical limitations, including insufficient chamber designs for hydrated samples and the lack of validated strategies for analyzing cellular O-PTIR data. The authors identify several persistent challenges associated with O-PTIR measurements in water. These include insufficient adaptation of measurement chamber designs for reliable positioning and stabilization of hydrated samples, as well as difficulties in maintaining accurate focus on cells with limited optical contrast when imaged using air objectives. In addition, signal instability during measurements in aqueous environments, manifested as fluctuations in DC images, further complicates reliable data acquisition. Here we aim to address the fundamental question of how to measure cells in an unaffected manner using O-PTIR spectroscopy and how to extract useful information from the collected data. A further consideration was how the O-PTIR data can complement information derived from RS, considering the latter is of high resolution and can accurately characterize cellular structures, especially lipids, in an aqueous environment.

## 2. Materials and Methods

### 2.1. Sample preparation

HeLa cells (HeLa Human cell line, MERCK, cat. no 93021013-1VL), were cultured in Medium Essential Medium (MEM, Sigma Aldrich, cat. no M7145-100ML) supplemented with 1% L-Glutamine solutions (Sigma Aldrich, cat. no G7513-100ML), 1% MEM Non-essential Amino Acid Solution (Sigma Aldrich, cat. no M7145-100ML), 1% antibiotics (Sigma Aldrich, cat. no P4333) and 10% Fetal Bovine Serum (FBS, ThermoFisher, cat. no A5256701). Cell cultures were maintained at 37 °C in a 5 % CO2 atmosphere, passaged every 3 to 4 days when 80 % confluency was reached. HeLa cells were seeded on sterilized CaF_2_ slides with a diameter of 10 mm and 0.35 mm thickness (Crystran, England) at a density of 20,000 cells per well in a 24-well plate and left for 24 hours to adhere. On the day of treatment, the culture medium was replaced with a fresh medium. The d_34_-Oleic Acid (d_34_-OA, Sigma Aldrich) solution was added directly to the medium at 200 µM and 10 µM concentrations. To facilitate fatty acid (FAs) transport through the cell membrane, d_34_-OA was first saponified using NaOH (Sigma Aldrich) at 70 °C and then conjugated with bovine serum albumin (BSA, fatty acids-free, Sigma Aldrich). After 24h-incubation, cells were gently rinsed with Dulbecco’s Phosphate Buffer Solution (DPBS) and fixed for five minutes at room temperature with 2.5% glutaraldehyde (Sigma Aldrich). The fixed cells were kept in DPBS at 4°C until measurements were taken. Just before measurements, the cells were rinsed with room-temperature DPBS.

### 2.2. Raman measurements using a system for correlative measurements

Raman imaging of HeLa cells was performed using a confocal Raman microscope, WITec Alpha 300 (WITec GmbH, Ulm, Germany), equipped with air-cooled solid-state laser operating at 532 nm, along with a CCD detector (Andor Technology Ltd, Belfast, Northern Ireland). The laser power was equal to approximately 30 mW (measured before the objective). The RS spectra were obtained using a 60x water immersion objective (Nikon Fluor, NA=1) with an integration time of 0.5 s and step size of 0.5 μm. Data processing was conducted using WITec Project Plus software. HeLa cells were seeded on glass slides and marked with a sticker featuring a characteristic symbol with guidelines enabling precise sample positioning along the x and y axes. Using a fixed reference point allowed for precisely identifying the same cell across different platforms, ensuring accurate comparison of the data obtained. The calculations were performed using a Correscopy software (Correscopy™, Frysztag, Poland).

### 2.3. O-PTIR measurements

O-PTIR measurements were acquired with a mIRage-LS microscope (Photothermal Spectroscopy Corporation, Santa Barbara, CA, USA) equipped with a probe laser with an emission wavelength of 532 nm and a tunable pump Quantum Cascade Laser (QCL, Daylight Solution, US) operating in the ranges of 2700-3000 cm^-1^, 2000-2300 cm^-1^, and 900-1800 cm^-1^. All measurements were performed in a co-propagating geometry. Dry samples were measured using a 40x Cassegrain objective (PIKE Technologies, Fitchburg, WI, NA=0.78) for 532 nm and IR. The O-PTIR spectra and single-band images were acquired using 11% of the IR laser power, 4% of the probe laser power, with a spectra recipe to maximize SNR (2 cm^-1^ / pt 200 cm^-1^/s), and 10x detector gain. Single-band images and spectra of dried cells were collected with a pixel size of 0.3 μm. Detection was performed in probe light reflection mode (532 nm). Aqueous samples were prepared in a sandwich configuration consisting of a thin microscopic slide and a CaF_2_ substrate to keep the cells hydrated in a microvolume of PBS. Transmission illumination was used to facilitate visible focus on the sample. Hydrated HeLas were measured using a 40x Cassegrain objective in a sandwich. The O-PTIR spectra and single-band images were acquired at 11% of the IR laser power, 4% of the probe laser power, using a high SNR recipe, and 2x the detector gain. The signal was collected in transmission mode, which refers to the detection of the transmitted probe beam (532nm). Single-band images of water samples were collected with a pixel size of 0.3 μm.

### 2.4. O-PTIR spectra and image analysis

All O-PTIR data were processed and analyzed using PTIR Studio 4.6 software (Photothermal Spectroscopy Corporation (PSC), Santa Barbara, CA, USA). The spectrum pre-processing included smoothing (Savitzky-Golay algorithm, 3^rd^ polynomial order, 11 points), normalization to the amide II band, calculation of the second derivatives (averaging width equal to 20 points), and a subsequent normalization to the amide II band. Normalization was performed in the amide II region because a strong background elevation due to water affects the most intense amide I band. The acquired single-channel images were displayed using a color-coded scale chosen based on the image histogram, aiming to highlight best the analyzed areas of the cell (LDs, nucleus, and complete cellular structure). Image analysis included calculating ratios for selected images and performing RGB analysis by overlaying chosen image layers. To improve the quality of images with very low values after calculations of ratiometric images, smoothing was applied (5 points).

## 3. Results and Discussion

### 3.1. Uptake of unsaturated FAs monitored by O-PTIR

We employed O-PTIR imaging to investigate the uptake of exogenous FAs by cancer cells, serving both as a model system to showcase the technique’s capabilities and to discuss the limitations of different approaches to analyze intracellular biochemical processes. To investigate the incorporation of unsaturated FAs, HeLa cells were incubated with d_34_-OA, for which the signals from C-D stretching vibrations are visible in the spectral silent region (1800-2700 cm^-1^), where there are no signals from cellular components. This approach allows for reliable monitoring of the uptake and intracellular distribution of FAs, facilitating the separation of signals from endogenous and exogenous substances.

As the first step, we used a common approach involving fixed and dried cells, a widely adopted strategy in IR imaging. Single-channel O-PTIR imaging was performed at selected wavenumbers, characterizing vibrations of biomolecules commonly found in cells, i.e. 1245 cm^-1^ (ν _as_(PO_2_^-^) in nucleic acids and phospholipids), 1660 cm^-1^, 1550 cm^-1^ (amide I and amide II (ν (C=O), δ(N-H) and ν (C-N)) in proteins), 1745 cm^-1^ (ν (C=O) in triacylglycerols (TAGs)), and 2100 cm^-1^ (ν _as_(C-D) in d_34_-OA) (**Figure 1**). The selected wavenumbers were chosen for their uniqueness in characterizing specific subcellular regions, allowing for pinpointing areas of interest, such as those rich in lipids in LDs, or the nuclear compartment, thus providing a detailed view of cellular structures, as presented in **Figure 1**.

**Fig. 1.**
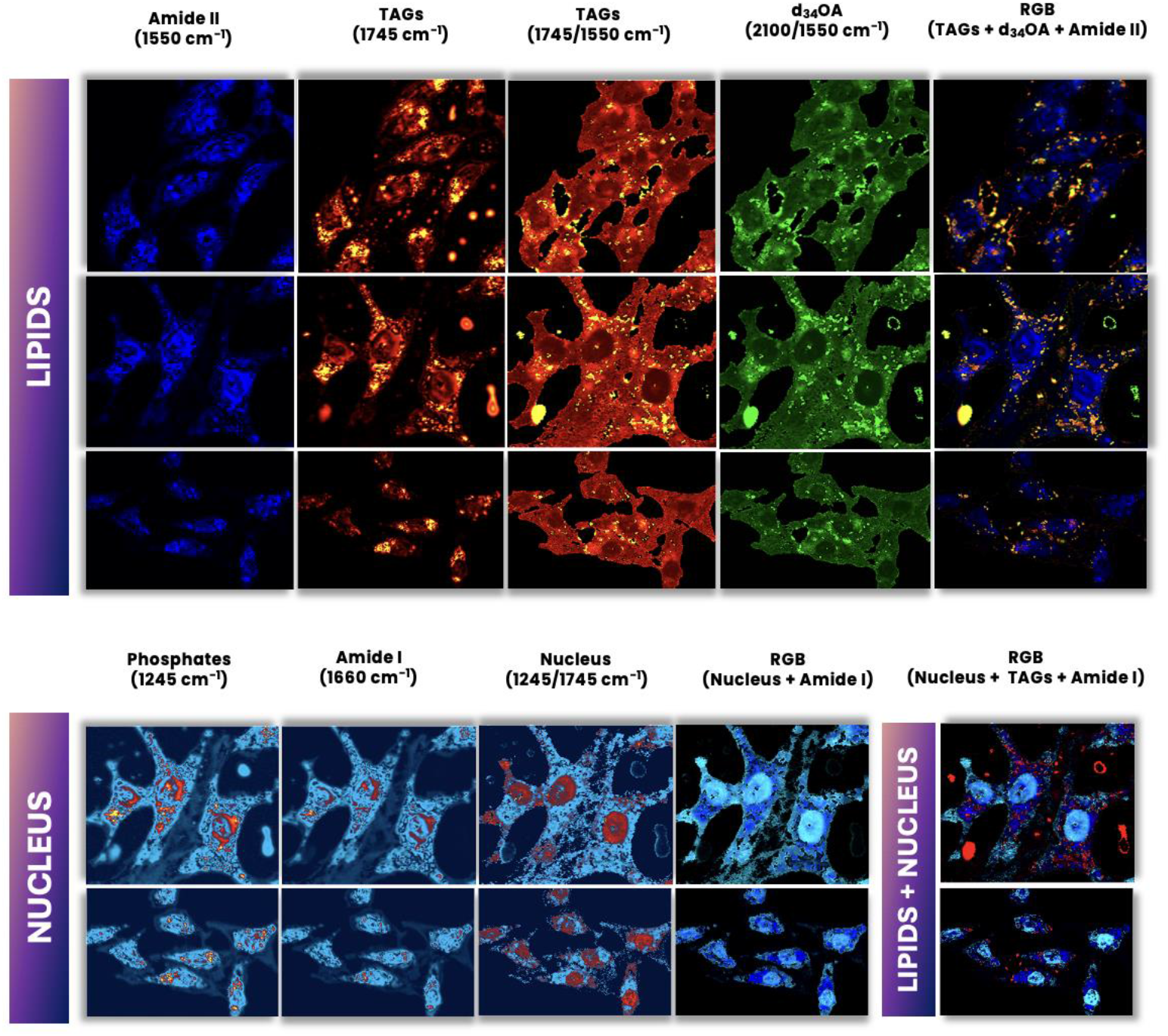
Single-band O-PTIR images of fixed and dried HeLa cells incubated with 200 µM d_34_-OA for 24h, showing the localization of subcellular regions enriched in lipids containing TAGs (1745 cm^-1^) and d_34_-OA (2100 cm^-1^), as well as the nuclear compartment (1245 cm^-1^). Regions of interest within cellular structures were highlighted using image ratio analysis and RGB overlay. The O-PTIR images acquired at different wavenumbers were presented in different color scales: 1245 cm^-1^ (0-20 mV), 1550 cm^-1^ (0 – 20 mV), 1660 cm^-1^ (0-20 mV), 1745 cm^-1^ (0-10 mV), 1745/1550 cm^-1^ (0-1 mV), 2100/1550 cm^-1^ (0-0.35 mV), and 1245/1745 cm^-1^ (0-7 mV).

While collecting chemical images at individual wavenumbers provides information about the distribution of specific biomolecules occurring in organelles, the interpretation of the obtained results is not always straightforward (**Figure 1**). In O-PTIR, the signal intensity generally exhibits a complex relationship with the size of the measured object, as it is influenced not only by the peak temperature but also by the dynamics of thermal decay, affecting the photothermal signal detected by the probe beam^28^. Therefore, it should be remembered that, firstly, there is a relationship between the size of the measured sample and the intensity of the detected O-PTIR signal, and secondly, this relationship may exhibit nonlinear behavior. To obtain reliable chemical information while minimizing the influence of sample thickness, a practical approach is to perform ratiometric analysis, image subtraction, or apply RGB overlays.

For example, calculating the ratio of 1745 cm^-1^ to 1550 cm^-1^ enables the identification of changes in the accumulation of TAGs related to the formation of LDs, because it normalizes the lipid-related signal (ν (C=O)) to the protein-related signal (amide II), which depends on the cell thickness. The 1745/1555 cm^-1^ ratiometric image highlights areas in cytoplasm with increased lipid accumulation more effectively than single-wavenumber images alone (**Figure 1**). Ratiometric images exhibit better contrast and chemical specificity, enabling the precise location of LDs. Similarly, calculating the 1245/1745 cm^-1^ ratio improves the visualization of the nuclear compartment. This is because the signal at 1245 cm^-1^, which is mainly associated with nucleic acids abundant in the nucleus, was normalized to the lipid signal at 1745 cm^-1^. As a result, the ratiometric image highlights areas of high nucleic acid-to-lipid signal, effectively delineating the nucleus from lipid-rich cytoplasmic regions.

Calculating ratiometric imaging facilitates chemical analysis but requires the selection of images of appropriate quality and careful interpretation of the obtained results. Calculating ratios in regions of low signal intensity, such as areas containing biological debris or thin cell membranes at the edges of the cell, can lead to artificially high ratio values, resulting in visual artifacts (**Figures S1 and S2**). Similarly, this is relevant when determining the ratio of background signals. Additionally, dividing images collected at wavenumbers related to bands with significant absolute intensity differences can amplify noise or introduce distortions, particularly when one of the signals approaches the detection limit. Therefore, prior assessment of the spectral profile of samples that will be further imaged, as well as maintaining an adequate signal-to-noise ratio (SNR) in the spectral images, is essential for reliable data interpretation^29^.

O-PTIR-based visualization of specific cellular organelles can be further improved through histogram analysis, providing information about the variability of the spectral signal across different images and their specific regions. Choosing the optimal range of the color scale using histograms and signal thresholding reduces potential subjectivity in image analysis^30,31^, as presented in the **Supporting Information** (SI). **Figures S1 and S2** illustrate how different scale selections can result in the loss of cellular features, such as membrane connections between cells, in single-channel O-PTIR images or ratiometric images, respectively. Ensuring a consistent dynamic range of O-PTIR signal intensities across images enhances the reliability of organelle identification and the evaluation of their spatial distribution.

Chemical images indicating the spatial distribution of individual biomolecules are valuable for analyzing the biochemical profile of cells. However, the true analytical power lies in multispectral imaging that provides morphochemical information across multiple biochemical features within the same cellular region simultaneously, offering a more comprehensive, multidimensional insight into subcellular organization. This is achieved through RGB overlays, yielding multi-channel O-PTIR images. The superimposition of 2100 cm^-1^ /1555 cm^-1^ and 1745 cm^-1^ /1555 cm^-1^ ratiometric images highlights areas of exogenous FAs accumulation together with LDs formation, respectively (**Figure 1**). This enables the distinction between endogenous and exogenous lipid pools within the cell. Additionally, incorporating the 1245 cm^-1^ / 1745 cm^1^ ratiometric image further enables the visualization of the cell nucleus.

Combining O-PTIR imaging with the proposed spectroscopic data processing methods is a practical approach that may provide an alternative to conventional cell staining techniques such as labelled fluorescence microscopy. By utilizing the intrinsic vibrational signatures of cellular components and enhancing their visualization through operations like ratiometric analysis and RGB overlays, it becomes possible to visualize and distinguish subcellular structures with high specificity. This strategy reduces sample preparation complexity and preserves the natural biochemical environment, making it a powerful tool for non-invasive cellular imaging. However, to fully utilize the potential of IR spectroscopic imaging for cells, it is necessary to use the measurement method of cells in an aqueous environment.

### 3.2. A Robust “Cell-In-Water” Approach

It is well established that cell dehydration disrupts the delicate cellular architecture by altering the integrity of the cell membrane, collapsing the cytoskeleton, and causing shrinkage or deformation of organelles^32^. Therefore, optimizing IR imaging techniques for analyzing cells in their native, hydrated state is essential. In this paper, we improved sample preparation to perform subcellular chemical analysis of HeLa cells and monitor the uptake of d_34_-OA under physiological conditions using O-PTIR. We also performed comparative O-PTIR measurements of the same single cells in buffer and the dehydrated state (**Figure 2**) to verify how drying affects their biochemical structure and morphology. For measurements in buffer, cells were enclosed in a sandwich configuration to maintain a physiological environment during the measurements, allowing for prolonged hydration. Sealing of the sample edges with two-component prosthetic silicone enabled the hydrated microenvironment to be maintained for up to 5 hours. The relatively long duration allows for spectral acquisition and imaging at single wavenumbers, as well as hyperspectral imaging. In aqueous conditions, measurements of adherent cells on glass substrates are particularly challenging due to their optical transparency, which complicates brightfield focusing and requires data collection in transmission mode, which can affect signal quality. Therefore, all measurements were performed using an optimal configuration, i.e., co-propagation geometry with transmission detection of the probe beam, which allowed for the acquisition of both images and single cell spectra with subcellular spatial resolution (**Figure 2A**). The proposed strategy of sample preparation and signal acquisition ensures stable measurement conditions for cells in buffer for an extended period while minimizing the signal contribution from extracellular water due to the small volume in the measurement chamber (∼ 2 µl). To the best of authors’ knowledge, this methodology currently represents the most effective approach for performing O-PTIR measurements of cells in an aqueous environment. While prior studies of O-PTIR measurements in water have largely emphasized spectral results, imaging data—particularly DC images—are rarely reported. DC images represent the average probe beam intensity scattered from the sample and provide bright-field–like contrast. In this work, DC images are included (SI) to enable direct comparison between dry and hydrated measurements, thereby illustrating the reduced stability commonly observed during O-PTIR imaging in aqueous environments. Following evaluation of the proposed methodology, DC images obtained using this approach exhibit the highest sharpness and contrast among the tested configurations.

**Fig. 2.**
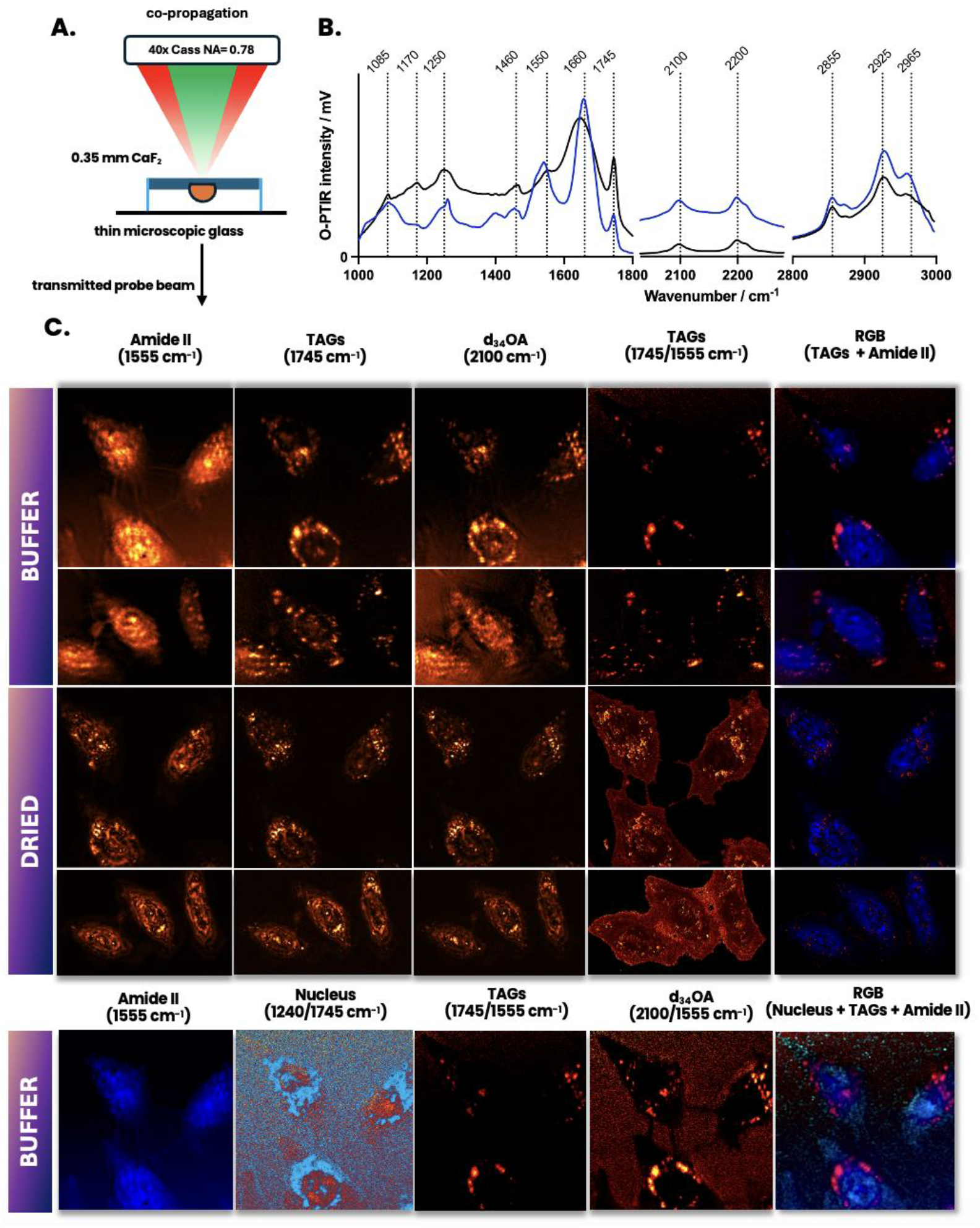
**A**. Schematic representation of the ‘sandwich’ configuration. **B**. Average O-PTIR spectra from LDs under hydrated (black) and dehydrated conditions (blue). The spectra were recorded from approximately 20 cells *per* sample. **C**. O-PTIR images showing LDs distribution in cells kept in buffer microenvironment (top) and after drying (bottom). The same cellular areas were analyzed for both conditions to facilitate a better comparison. Before measurements, HeLa cells were incubated with 10 µM d_34_-OA for 24 h. RGB Overlay represents the superimposition of ratiometric images: 1745/1555 cm^-1^, 1240/1745 cm^-1^, and the image collected at 1555 cm^-1^.

Using the developed methodology, cells were imaged in the hydrated state, and then spectra from the selected regions of interest were acquired. **Figure 2B** shows a comparison of the average LDs spectra from dry and wet cells, illustrating how the spectrum changes after dehydration in comparison to aqueous conditions. The spectrum from the cell in buffer exhibited significant background elevation and band broadening in the silent and fingerprint regions in comparison to the spectrum collected from dried cells. This is particularly evident in the amide I (1660 cm^-1^) and amide II (1550 cm^-1^) bands, which display increased full width at half maximum (FWHM), with the amide II band barely distinguishable in buffer. Second derivatives of the presented O-PTIR spectra were calculated to identify more subtle changes in their spectral features and are presented in **Figure S3** along with band assignments **(Table S1)**. Our results show that, due to the presence of water primarily within cells, water signals remain visible in O-PTIR spectra under hydrated conditions and can interfere with protein secondary structure analysis. Only precise focus on the sample and minimizing the water surrounding the cell can guarantee high-quality data acquisition with a reduced water background.

In O-PTIR imaging, water is a disturbance that increases the background signal around the cells. **Figure 2C** shows a comparison of images obtained from single cells and their analysis in wet and dry states. Single-cell measurements allowed comparison of the same cells in both conditions. As can be seen, the LD image in cells changes significantly before and after drying, as observed in images collected at 1745 cm^-1^ and 2100 cm^-1^. The lipid storage structures in dried cells appear more granular compared to hydrated cells. This may be due to the collapse of the cytoskeleton, which maintains the structural integrity of cells, and the degradation of subcellular structures. Therefore, the proper cellular organization of lipid-rich structures is likely disrupted under dried conditions. O-PTIR images of hydrated cells collected at 1555 cm^-1^ show a more uniform distribution of proteins within the cells, ensuring the maintenance of proper cellular morphology.

As with dried samples, normalization of the 1745 cm^-1^ images to the spectral images representing the protein distribution (amide II) demonstrated significantly better contrast in LD imaging in water than the single-wavenumber O-PTIR images. Additionally, ratiometric 2100/1555 cm^-1^ images accurately revealed d_34_-OA accumulation. It is also worth highlighting that in the case of measurements performed on cells in buffer, the incubation concentration was 20-fold lower (10 µM) than for dried cells presented in Figure 1. Nevertheless, the developed methodology still allowed for signal detection. This demonstrates the high sensitivity of this approach, despite some limitations related to the elevated background level. Similarly to what was observed for dried samples, histogram analysis and thresholding improve accuracy, which is particularly important when calculating ratiometric images. This minimizes high background signals or artifacts resulting from division by low-intensity values, as shown in **Figures S2 and S4**.

### 3.3. Complementarity of O-PTIR and RS in Lipid Analysis

The same hydrated cell imaged by O-PTIR was also measured in aqueous conditions using the RS technique (**Figure 3**), known for its high spatial resolution, ability to perform measurements in buffer, and high sensitivity to lipids^33^. Additionally, RS captures hyperspectral images, providing rich information about the biochemical landscape of cells and the distribution of their components^34^. This makes RS highly suitable for verifying whether the developed methodology for performing O-PTIR measurements in aqueous conditions effectively preserves cellular structure, as well as for assessing the complementarity of O-PTIR and RS in subcellular analysis under physiological conditions, using the example of d_34_-OA uptake in HeLa cells as a model system.

**Fig. 3.**
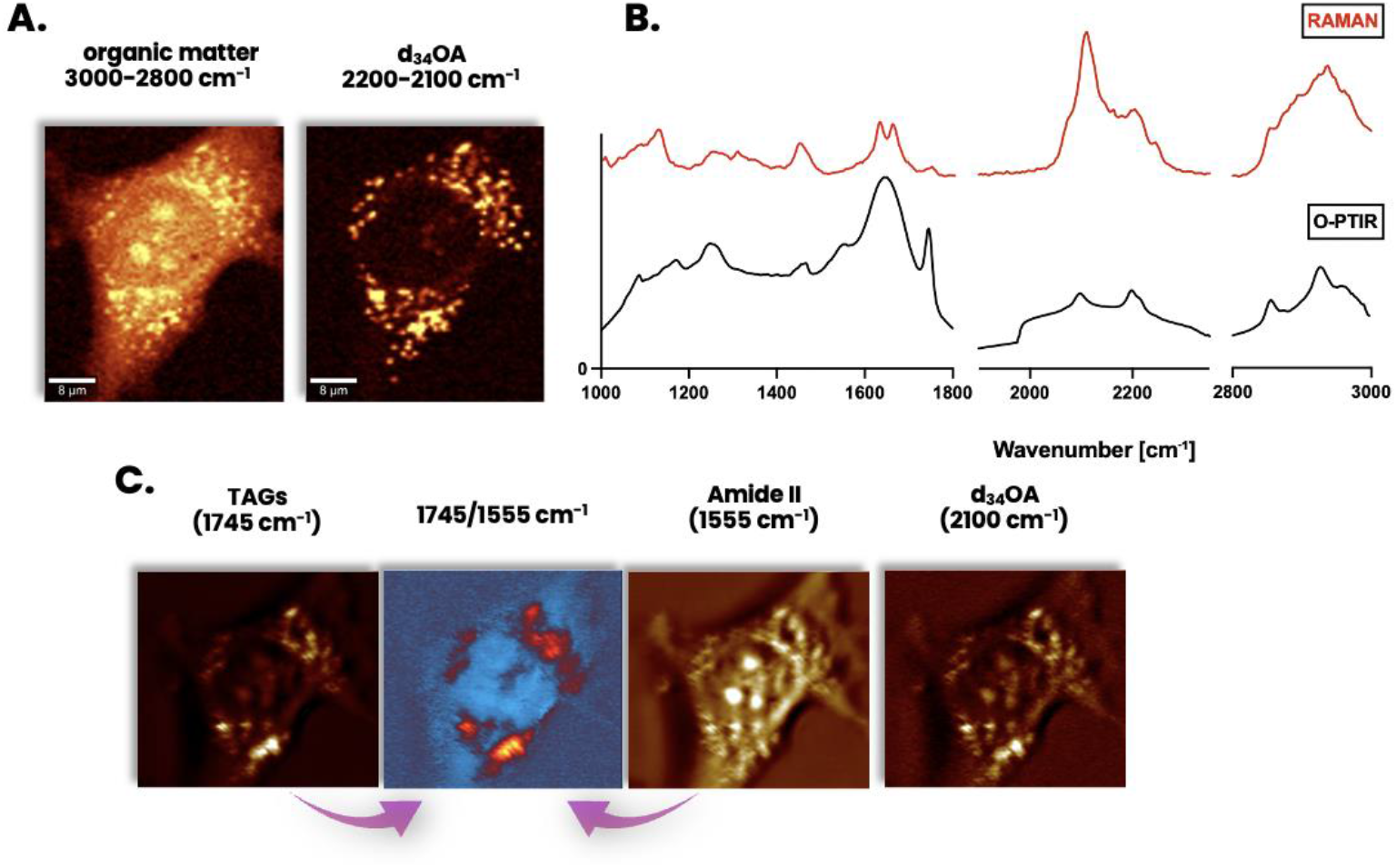
Comparison of RS and O-PTIR images of the same HeLa cell measured under aqueous conditions. **A**. The RS images were obtained by integration of bands typical for organic matter (2800-3000 cm^-1^) and d_34_-OA (2100-2200 cm^-1^). **B**. Comparison of RS (red) and O-PTIR (black) representative spectra from lipid-rich area within the single cell in buffer. **C**. The O-PTIR images were acquired at 1555 cm^-1^, 1745 cm^-1^, and 2100 cm^-1^ to compare the probe-specific signals within the single cell.

The distribution of exogenous FAs was visualized using RS through a chemical image obtained by integrating the d_34_-OA specific band (i.e., 2100–2200 cm^-1^). This was compared with the O-PTIR image collected at 2100 cm^-1^. Furthermore, a comparative analysis of the TAGs distribution in LD was conducted in the same manner using O-PTIR images (band analysis at 1745 cm^-1^ and the ratiometric image at 1745/1550 cm^-1^). Due to their low intensity, these bands were not integrated for the RS data. Cell morphology was visualized by integrating RS features corresponding to organic matter in the range of 2800–3000 cm^-1^ and compared with the protein distribution obtained by registering O-PTIR signals at 1550 cm^-1^ (amide II).

The results obtained using the RS and O-PTIR techniques were compared and showed a significant difference in image quality and resolution. RS images show even single LDs, whereas O-PTIR images only show entire clusters. This highlights the difficulties encountered when O-PTIR imaging in aqueous environments. Furthermore, analyzing the ratios of O-PTIR images leads to the loss of important information regarding the LDs’ location. A key advantage of RS in this context is the availability of water-immersion and confocal objectives, which facilitate improved focusing and spatial resolution in hydrated samples. Such optical configurations are not available in the standard Mirage-LS setup, and their implementation may represent an important future step toward enhancing O-PTIR imaging performance in aqueous conditions. Other problems are most likely due to the significant influence of the water background and the low signal stability during O-PTIR measurements in aqueous conditions, despite the use of an optimized measurement strategy.

However, spectral analysis revealed that O-PTIR offers complementary information, distinct from that obtained with RS. Bands in O-PTIR spectra other than those in RS spectra can be used to characterize biological material. After incubation of cells with d_34_-OA, it was observed that not only typical signals, specific to C-D or TAG vibrations were present in the O-PTIR spectrum, but also other signals, such as 1085 cm^-1^, 1250 cm^-1^, 1460 cm^-1^, and 1745 cm^-1^, strongly co-localize with the presence of LDs, showing for example, the distribution of phospholipids, and TAGs. The O-PTIR spectral profile differs from that observed in RS, where the characteristic lipid bands mainly include 1250 cm^-1^, 1305 cm^-1^, 1440 cm^-1^, 1660 cm^-1^, and 2850 cm^-1^, where phospholipids can be detected, but observation of TAGs is significantly more difficult. This highlights the complementarity of both methods and the need to combine techniques in routine studies, using, for example, data fusion approaches^35,36^. Only by combining the information obtained from both techniques can a detailed and comprehensive analysis of the molecular structure of the studied cells in hydrated environments be performed. O-PTIR and RS are characterised by differently sensitivities to the chemical composition of different types of biomolecules (such as proteins or lipids), and their combination, rather than the use of a single technique provides an effective approach in cell characterization^37,38^.

### 3.4. Summary

In this work, high-resolution O-PTIR imaging of HeLa cells was performed. The cells were incubated with deuterated fatty acid d_34_-OA. After applying various analytical approaches, such as ratiometric image analysis and histogram evaluation, reliable distribution maps of selected subcellular components were obtained. The distribution of exogenous fatty acids (FAs) and TAGs have been effectively visualized by normalizing the vibrational signals of C–D bonds (2100 cm^-1^) and vibrational signals of ester groups (1745 cm^-1^) to signals derived from proteins, such as those from amide bands (amide II, 1550 cm^-1^). This approach may be an alternative to fluorescence labeling microscopy, enabling the simultaneous visualization of several subcellular structures based on their intrinsic vibrational signatures.

A new reliable measurement protocol has been developed that enables the study of cells in their natural environment, i.e., in a buffer. While earlier studies have shown that O-PTIR and FT-IR measurements of cells in water are possible, they often do not provide sufficient guidance for overcoming critical challenges related to sample preparation, signal monitoring, and analysis of subcellular features. We propose placing cells in a measurement chamber sealed with silicone adhesive, which ensures stable measurement conditions in the buffer for up to 5 hours. This time is sufficient for both imaging and acquisition of single-cell spectra. Furthermore, the small volume of liquid in the measurement chamber (2 µl) minimizes the contribution of signals from extracellular water. Comparison of chemical images obtained using O-PTIR for the same cells under hydrated and dehydrated conditions, as well as a direct comparison of O-PTIR and RS, demonstrates that cellular structure is well preserved in an aqueous environment. The obtained data more accurately reflect the correct morphology and structural organization of cells. However, our results show that despite the development of an improved protocol, problems remain, such as elevated background signals, overlap with water-derived signals (including intracellular water), and instability of the O-PTIR signal. These problems can be partially mitigated through various image processing techniques, such as ratiometric analysis. However, further optimization of the measurement configurations is necessary to minimize the influence of instrumental and experimental factors on the obtained results. In summary, this study addresses practical challenges commonly encountered when performing O-PTIR measurements in aqueous environments. It highlights the importance of minimizing the sample volume (∼2 µL in a microchamber), monitoring signal stability through DC images, and implementing careful data analysis protocols. Collectively, these considerations provide a robust framework for reliable subcellular measurements in hydrated cells and offer guidance for future studies aiming to perform high-resolution infrared imaging under physiologically relevant conditions.

Comparison of RS and O-PTIR imaging of the same cell demonstrated that, at the subcellular level, combining these two spectroscopic modalities provides comprehensive and complementary information. RS and O-PTIR appear to be a harmonious combination that works effectively together, and further development of integrated approaches, including measurement strategies and analytical frameworks such as data fusion, is of immense value. Undoubtedly, such integrated strategies will provide deeper insights than ever before into the complex and intricate mechanisms operating at the subcellular level.

## Acknowledgments

This research has been funded by the National Science Center, Poland (NCN), Maestro 2022/46/A/ST4/00054 to Malgorzata Baranska. This publication has been funded by the program *Excellence Initiative – Research University* at the Jagiellonian University.

